# Rewiring of peatland plant-microbe networks outpaces species turnover

**DOI:** 10.1101/2020.05.12.090274

**Authors:** Bjorn J.M. Robroek, Magalí Martí, Bo H. Svensson, Marc G. Dumont, Annelies J. Veraart, Vincent E.J. Jassey

**Author notes:** These authors contributed equally to this work (Bjorn J.M. Robroek, Magalí Martí). Author for correspondence: B.J.M. Robroek.

## Abstract

Enviro-climatological changes are thought to be causing alterations in ecosystem processes through shifts in plant and microbial communities; however, how links between plant and microbial communities change with enviro-climatological change is likely to be less straightforward but may be fundamental for many ecological processes. To address this, we assessed the composition of the plant community and the prokaryotic community –using amplicon-based sequencing– of three European peatlands that were distinct in enviro-climatological conditions. Bipartite networks were used to construct site-specific plant-prokaryote co-occurrence networks. Our data show that between sites, plant and prokaryotic communities differ and that turnover in interactions between the communities was complex. Essentially, turnover in plant-microbial interactions is much faster than turnover in the respective communities. Our findings suggest that network rewiring does largely result from novel associations between species that are common and shared across the networks. Turnover in network composition is largely driven by novel interactions between a core community of plants and microorganisms. Taken together our results indicate that plant-microbe associations are context dependent, and that changes in enviro-climatological conditions will likely lead to network rewiring. Integrating turnover in plant-microbe interactions into studies that assess the impact of enviro-climatological change on peatland ecosystems is essential to understand ecosystem dynamics and must be combined with studies on the impact of these changes on ecosystem processes.

## Introduction

Throughout the present interglacial period, northern peatlands have acted as carbon (C) sinks, resulting in over 500 Gt of C (Yu et al. 2010) to be locked up in these ecosystems. Recently it has even been suggested that northern peatlands may contain twice as much C (Nichols & Peteet 2019), i.e. > 1000 Gt, yet these numbers are debated (Ratcliffe et al. 2020). Nevertheless, maintaining a positive carbon balance (C uptake > C release) in peatlands is of great relevance to mitigate carbon-climate feedbacks. The carbon sink function of peatlands largely depends on the imbalance between gross primary production and decomposition. In undisturbed peatlands, the balance of these two processes is largely in favour of primary production. Indeed, decomposition rates in natural peatlands are low. Peatland plants, and *Sphagnum* mosses in particular, produce recalcitrant litter (Clymo 1965, Dorrepaal et al. 2005) and release anti-microbial compounds (Fudyma et al. 2019, Hamard et al. 2019), which together with acidic and anoxic environmental conditions impede microbial breakdown of organic matter and thus facilitate the accumulation of plant remains. Hence, the C sink function of peatlands is controlled by an interplay between abiotic and biotic factors.

Direct effects of environmental change on peatland ecosystem function have been studied intensively. Examples of these direct effects include changes in nitrogen deposition (Aldous 2002, Bragazza et al. 2006, Olid et al 2014), drought (Fenner & Freeman 2011; Estop-Aragonés et al. 2016), and warming (Dorrepaal et al. 2009, Olid et al 2014, Wilson et al. 2016), amongst others, which have been linked to carbon loss. Environmental change can however also indirectly affect ecosystem function, through alterations in the biotic composition of ecosystems (Walther et al. 2002, Pimm et al. 2014). Such changes in community composition more often than not have consequences for ecosystem processes (Bardgett et al. 2013, Rillig et al. 2019). Indeed, the effects of warming, drought and nutrient deposition on carbon cycling in peatlands have previously shown to manifest via changes in the composition of the vegetation (Limpens et al. 2003, Dieleman et al. 2015, Rupp et al. 2019). However, the effect of environmental change on carbon cycling can also be mitigated by changes in the vegetation through plant-controlled metabolomic and biogeochemical mechanisms that protects C loss during drought (Wang et al. 2015; Fenner & Freeman, 2020). Following the train of thought that changes in environmental conditions cause shifts in the composition of peatland plant communities (Vitt & Slack 1975, Robroek et al. 2017, Norby et al. 2019), convergent shifts in the microbial community can be expected. Indeed, peatland microbial community structure and activity are strongly connected to plant community assemblage (Bragina et al. 2014, Chronakova et al. 2019, Martí et al. 2019b, Ivanova et al. 2020). Hence, interactions between plant and microorganisms are the cornerstone that shape carbon-related processes in peatlands (Lindo et al. 2013) and it is critical that we understand how enviro-climatological change affects these interactions to forecast the consequences for ecosystem processes (Kostka et al. 2016).

Most species do not exist in isolation but are embedded in trophic, non-trophic, mutualistic or antagonist interaction networks (Tylianakis et al. 2008, Bascompte 2009, Kéfi et al. 2016). In these networks, the number of species and interactions vary depending on species identity, functional traits present and enviro-climatologic niche (Schleuning et al. 2016). Network assemblage, in turn, can affect the resilience and stability of the network, which ultimately underlies the robustness of ecosystem functions to enviro-climatic change (Allesina et al. 2009, Morriën et al. 2017, de Vries et al. 2018). The effects of changes in plant community composition on soil microbial communities, are however thought to be dependent on the environmental context (De Vries et al. 2012) – but see Fanin, et al. (2019) –, leading to complex responses of plant-microbial interactions to changes in environmental conditions (Hagedorn et al. 2019). Adding to this complexity, recent work in peatlands has shown that plant and soil communities exhibit non-linear responses to changing environmental conditions (Jassey et al. 2018, Lamentowicz et al. 2019), yet no studies have tested the co-response of peatland plant and microbial communities to environmental change. To overcome this knowledge gap, it is essential to gain understanding on how environmental conditions regulate plant-microbe associations (i.e. networks) and how this relates to within-community responses (i.e. plant and microbial communities). Such knowledge would ultimately lead to improved understanding on the effects of environmental change on peatland processes.

To assess how enviro-climatological conditions affect plant-microbe interactions, we characterized plant and microbial communities in three European *Sphagnum-dominated* peatlands. We assessed how enviro-climatological conditions relate to compositional differences in plant and 16S rRNA-derived microbial communities (i.e. occurrence of taxonomic units with active function) and how this affects plant-microbial networks, including network topology. We postulate that enviro-climatic change will trigger both communities to turnover, leading to changes in the network structure. The influence of enviro-climatologic conditions on plant-microbe associations can manifest in various ways (Fig. 1): (i) no difference in biotic communities, nor in network associations, (ii) no difference in biotic communities but a rewiring in network association with altered enviro-climatological conditions, (iii) differences in the composition in one or both communities with congruent differences in network associations, and (iv) differences in biotic community compositions with a disproportionate rewiring of interaction networks. Congruent with the effects of environmental conditions on peatland plant communities (Robroek et al. 2017), we expect that differences in plant-microbe network structure are largely driven by turnover in one or both communities.

**Figure 1.**
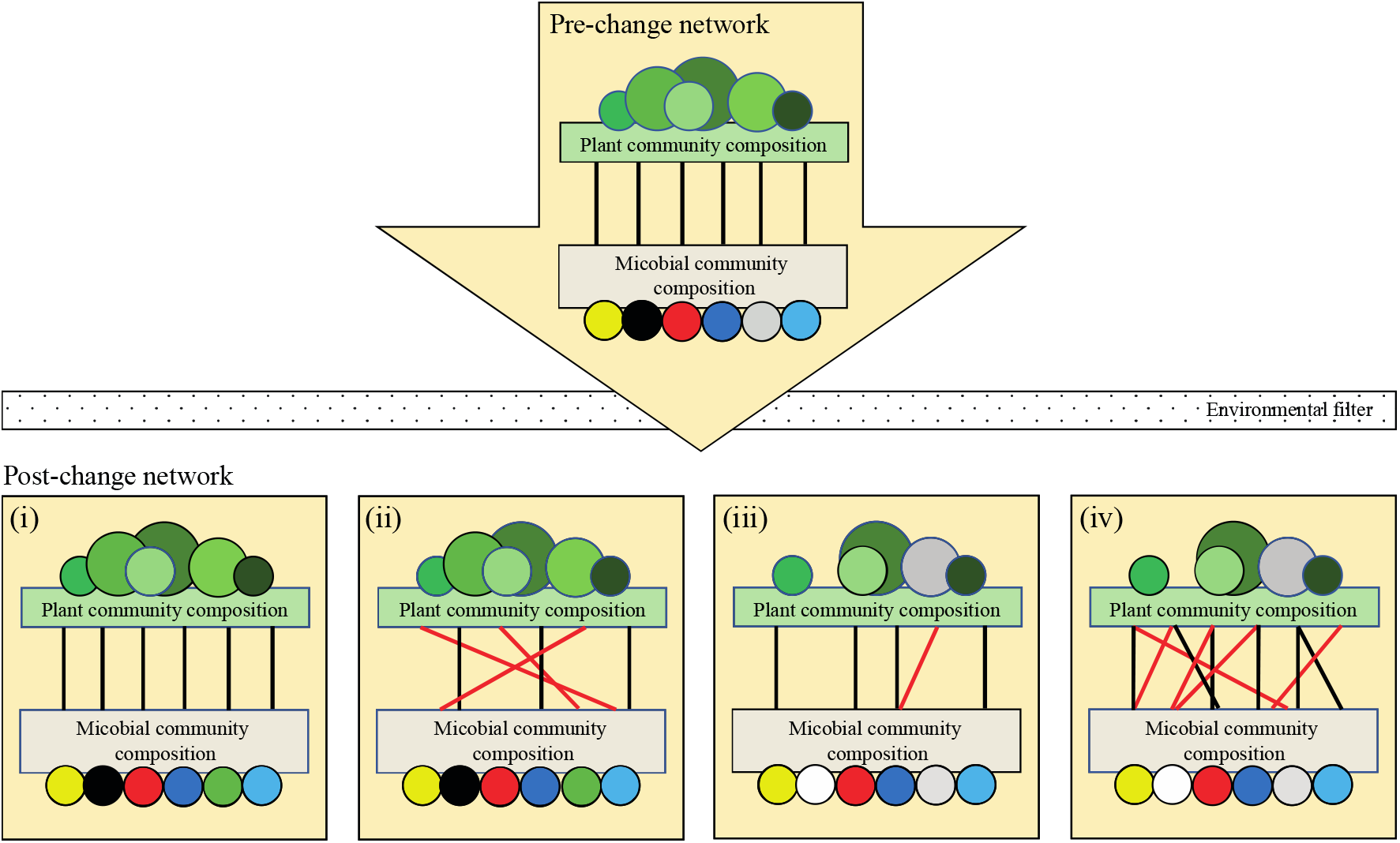
Examples of potential, i.e. hypothesised, effects of enviro-climatological change on associations between species in the plant community and ‘species’ in the microbial community: plant-microbe networks. Effects of enviro-climatological change (environmental filter) on network structure (pre-change network) are postulated to manifest in four potential ways: (i) no alterations in post-change biotic communities, nor in network associations; (ii) no alterations in biotic communities but a rewiring in network associations [depicted as red lines]; (iii) alterations in the composition in one or both communities [depicted as changes in composition of coloured symbols] with congruent alterations in network associations, and (iv) alterations in biotic community compositions with a disproportionate rewiring of interaction networks.

## Material and Methods

### Site description and enviro-climatic data

Three geographically distinct European *Sphagnum*-dominated peatlands were selected for this study. Degerö Stormyr (64°88’N, 19°56’E, 277 m above sea level (asl)), Sweden, is a minerogenic peatland with ombrotrophic elements. Cena Mire (56°25’N, 23°81’E, 12 m asl), Latvia, is an ombrotrophic raised bog. Dosenmoor, (54°95’N, 10°02’E, 28 m asl), Germany, is a restored ombrotrophic raised bog. These sites vary in degree of conservation status, but all sampling locations were located in an ombrotrophic part of each peatland with vegetation characteristic for natural peatlands. Full vegetation characteristics of the sites can be found on https://doi.org/10.5061/dryad.g1pk3.

We collected variables that describe climatic and environmental conditions in the selected peatlands previously being shown to be the most important drivers for the turnover in peatland plant community composition (Robroek et al. 2017). Four bioclimatic variables (mean annual temperature, temperature seasonality, mean annual precipitation, precipitation seasonality) were extracted from the *WorldClim* database (Hijmans et al. 2005), and averaged over a 5-year (2005-2009) period that preceded our sampling campaign. *WorldClim* (http://www.worldclim.org/bioclim) calculates temperature seasonality as the standard deviation of the mean monthly temperature. Larger standard deviations refer to a larger variability in monthly temperature. Precipitation seasonality is a measure of the variation in monthly precipitation totals and calculated as the standard deviation of the monthly precipitation estimates expressed as a percentage of the mean of those estimates (i.e. the annual mean). Atmospheric deposition data were produced using the EMEP (European Monitoring and Evaluation Programme)-based IDEM (Integrated Deposition Model) model (Pieterse et al. 2007) and consisted of grid cell (50 × 50 km) averages of total nitrogen and sulphur deposition. pH was measured (Mettler Toledo MP220 pH meter) in five wells in each peatland site. These wells were situated on the lawn-hummock interface.

### Field sampling of plant and microbial communities

Abundance data for all vascular plant and bryophyte species from randomly selected lawn (n =3) and hummock (n = 3) microhabitats (0.25 m^2^ quadrats; six in total) were collected in the summer of 2010. Microhabitats were sampled in pairs, with each lawn-hummock pair separated < 2 m. This sampling strategy was chosen as to represent the nature of the ombrotrophic part of the peatland as much as possible. Vascular plants and *Sphagnum* mosses were identified to the species level. *Non-Sphagnum* bryophytes were identified to the family level. Rarefaction analysis indicated that our sampling adequately captured species richness in our peatlands.

In all plots (3 sites, 2 microhabitats, 3 replicates; 18 in total), we collected a 5 cm^3^ peat sample from the acrotelm – defined as the peat layer between the peatland surface and the lowest water table level, which largely contains living plants – and the catotelm – the peat layer that underlies the acrotelm, is largely anoxic, and contains mostly dead plant remains (Clymo 1984). Hence, we collected a total of 36 samples from 18 plots. Sampling depth for the acrotelm samples was 5 cm below the surface, while sampling depth of the catotelm samples was 12.5 cm and 37.5 cm below the surface for the lawns and hummocks, respectively (Marti et al. 2015; Suppl. Fig. 2). Sampling was conducted using a 10-cm wide Holmen auger, which was carefully cleaned with 70% alcohol between sampling plots. Samples were refrigerated at 4°C, transported to the laboratory, and stored at −20°C prior to RNA extraction.

### RNA isolation, amplicon sequencing and data handling

Total RNA was extracted from 0.75 g of wet peat (7.8% ± 2.5 dry peat) using the FastRNA®Pro Soil-Direct Kit and FastPrep® Instrument (MP Biomedicals, Santa Ana, CA) as per the manufacturer’s instructions. Quality and concentration were determined on a Qubit fluorometer (Invitrogen, Eugene, OR, USA). Extracted RNA was reverse transcribed to complementary DNA (cDNA) using the lllustraTM Ready-to-Go RT-PCR Bead (GE-Healthcare, Uppsala, Sweden). Briefly, 2 μl of the extracted RNA and 2.5 μl of the random primer pd(N)6 (1.0 μg/μl) (GE-Healthcare, Uppsala, Sweden) were mixed, denatured at 97°C for 5 min, and then chilled on ice. One IllustraTM Ready-to-Go RT-PCR Bead and RNAse-free water were added to a final volume of 50 μl. Next, reverse transcription, consisting of a chain of 30 mins at 42°C followed by 5 mins at 95°C was used to create cDNA. RNA and cDNA were stored at −80°C and −20°C, respectively.

A 485 base pair (bp) fragment flanking the V3-V4 region of prokaryotic 16S rRNA was amplified using the primers 341F (5’-CCTAYGGGRBGCASCAG-3’) and 806R (5’-GGACTACNNGGGTATCTAAT-3’) modified from (Yu et al. 2005). Each 25-μl PCR mixture contained one Illustra PuReTaq Ready-To-Go PCR Bead (GE Healthcare, Uppsala, Sweden), 0.5 μM of each primer, 5 μl of cDNA and RNAse-free water. The 35 PCR cycles, performed on a MyCycler™ Thermal Cycler (Bio-Rad Laboratories), consisted of 90 s at 95°C, 30 s at 95°C, 30 s at 56°C, 30 s at 72°C and a final 5 mins at 72°C. PCR products were verified on 1% agarose gel and purified with a GeneJET PCR Purification Kit (Fermentas, Vilnius, Lithuania). Purified PCR product concentrations were determined using the Quant-iT dsDNA HS Assay Kit and the Qubit fluorometer (Invitrogen) and adjusted to 5 ng μl^-1^. To link sequence reads to samples, a second PCR on the initial product was performed using the primer pair 341F/806R with adapters and a unique barcode. The number of cycles was reduced to 10. Bands of ca. 530 bp were cut out and purified by High Pure PCR Cleanup Micro Kit (Roche Diagnostics GmbH, Mannheim, Germany) as per the manufacturer’s instructions. These fragments were quantified using a Qubit™ dsDNA HS Assay Kit and the Qubit™ fluorometer (Invitrogen) and qPCR (Mx-3000, Stratagene). The sample amplicons were mixed to equal concentrations (4 × 10^5^ copies μl^-1^) and subjected to two-region 454-pyrosequencing on a 70 × 75 GS PicoTiterPlate (PTP) using a GS FLX pyrosequencing system.

Sequence data were sorted, trimmed, filtered and quality checked (QS ≥ 25 for trimming, and lengths ≥ 150 bp) using the Qiime 1.6.0 data analysis pipeline (Caporaso et al. 2010). Chimeras were detected and removed using UCHIME in reference mode with the MicrobesOnline gold set (Edgar et al. 2011). Briefly, operational taxonomical units (OTUs) were selected at 97% identity with a confidence threshold of 50%. Representative sequences were selected randomly and classified against the SILVA database version 132 (https://www.arb-silva.de/). Unifrac distances for beta diversity were calculated from trees generated with Fasttree. The input multiple sequence alignments were generated using PyNAST against gg2011 template alignments. From the 36 libraries, we obtained a total of 772,742 sequences with an average of 21,465 sequences per library, except for one sample from Cena Mire –lawn, acrotelm– which had 135 reads and was therefore removed from the analysis. OTUs that were present in fewer than 20% of the samples were removed from the analyses as well as OTUs identified as chloroplast. A total of 464,327 sequences were left, corresponding to 4707 OTUs. From those, 74% (3690) of the OTUs corresponded to bacterial taxa and 20% (141) to archaeal taxa.6% of the OTUs were unclassified.

### Plant-prokaryote bipartite networks

Bipartite networks (i.e. a network with two distinct sets of nodes, plant nodes and prokaryotic nodes, with associations only defined between nodes from different sets) were used to detect co-occurrence between plants and prokaryotes as a proxy for sitespecific plant-prokaryote associations (Faust & Raes 2012). Co-occurrence networks were computed at site level. Hence data for the hummock and lawn microhabitats were pooled; however, separate networks were constructed to link the plant community with the acrotelm and with the catotelm prokaryote communities. Hellinger-transformed matrices, comprising data of bacterial and archaeal OTUs, vascular plants and *Sphagnum* species, were used to calculate Spearman’s correlation coefficients for plant-prokaryote associations (Berry & Widder 2014). Significance for each permutation was assessed using a permutation test with 10,000 repetitions. Only correlations > 0.6 (positive and negative) were adopted in the co-occurrence matrix (Sander et al. 2017). Each cooccurrence matrix was transformed into an adjacency (binary) matrix based on the presence or absence of links. Networks were then produced using the *igraph* and *bipartite* R packages (references in Supplementary), where each network comprised positive and negative plant-prokaryote associations. Each network was further analysed in terms of topology, and network dissimilarity (Berry & Widder 2014, Williams et al. 2014).

First, we evaluated the topological attributes of the networks using a set of indices focussed on species interactions, which were calculated for interaction between the plant and prokaryotic communities. *Effective partners* was defined as the number average of links for individuals in a specific network. *Partner diversity* is the diversity (PD; Shannon-based) of interactions between species from the two levels of the network. *Partner specificity* (PS) was calculated as the coefficient in variation of interactions, normalised to values between 0 and 1 (Poisot et al. 2012b), where PS = 0 indicates low specificity and PS = 1 indicates high specificity. These three indices were calculated from a top-down and bottom up perspective. Top-down accounts for the number of interactions plant species have with prokaryotic ‘species’. Bottom-up accounts for the number of interactions prokaryotes have with species in the plant community. Essentially, in a network top-down and bottom up networks do not necessarily have to match (Suppl. Fig. 3). Next, we identified network attributes using a set of plant-prokaryote interaction indices. *Linkage density* (LD) is the marginal total-weighted diversity of interactions between species. *Generality* of the plant community (G_p_) is calculated as the mean effective number of prokaryotic ‘species’ per plant species weighted by the total number of interactions. It describes the degree of specialization in the plant community. Low values indicate a high degree of specialization. *Generality* for the prokaryotic community (G_m_) is calculated as the mean effective number of plant species per prokaryotic ‘species’ weighted by the total number of interactions, and represents the degree of specialization in the prokaryotic community. *Robustness* (R) is a measure of robustness of the network to species loss, calculated as the area under the extinction curve of the network. The curve, essentially, describes the dependency –for survival– of species from one level on species of another level. R = 1 corresponds to a situation where most prokaryotes survive while most plant species are eliminated, or vice versa. R = 0 corresponds to a situation where species loss form one level results in the loss of most to all network connections. *Partner diversity* in the networks (PDn) is the diversity (Shannon-based) of the number of interactions for the species in each level of the bipartite network. *Species richness* (SR) is the number of species in the bipartite network community. *Niche specialization* (N) represents the site scores from a NMDS analysis, and represent the niche characteristics of each community (Devictor et al. 2010). The indices G, R, PD SR and N were calculated separately for the two levels –plants and prokaryotes– in the bipartite networks.

### Network turnover

To test if enviro-climatic conditions affected species turnover and turnover in bipartite networks, we calculated pairwise *β*-diversity in species composition and plant-prokaryote networks in each site using the methods proposed by Legendre & De Cáceres (2013) and Poisot et al. (2012a), respectively. For pairwise *β*-diversity in species composition, we used a site × species abundance matrix. Total *β*-diversity (BD) was partitioned into species contributions (BD_S_; degree of variation of individual species in each site) and local contribution (BD_L_; comparative indicators of the ecological uniqueness of the sites) to *β*-diversity (Legendre & De Cáceres 2013). For *β*-diversity in co-occurrence networks, we use a site × species pairs matrix (adjacency binary matrix). As differences in networks can arise either through changes in species composition and/or realized interactions between species, we specifically focused on the dissimilarities in interactions between networks (BDN) that originates from differences in species interactions due to species turnover (BDN_NS_) or from novel interactions between common network species (BDN_CS_) (Koleff et al. 2003, Canard et al. 2012, Poisot et al. 2012a).

### Data handling and statistical analyses

All statistical analyses were performed in *R* version 3.1.2 using the *vegan, diverse, phyloseq, nlme, dplyr and tidyr* packages (references in Supplementary). Analysis of Variance tests were used to test for differences in pore water nutrients, plant species and prokaryote richness, diversity (Shannon H’), evenness (Pielou’s J), and plant-prokaryote network indices between sites and sampling depth (for prokaryote data only). Prior to testing for turnover in prokaryotic community composition across the different habitats (β-diversity), relative abundance of each OTU was calculated within each sample.

We used redundancy analysis (RDA) to relate enviro-climatologic factors to plant and prokaryotic communities (Hellinger-transformed matrices). We used *ClustOfVar* package in R to reduce the enviro-climatological covariates of interest (i.e. mean annual temperature, temperature seasonality, mean annual precipitation, precipitation seasonality, total nitrogen and sulphur depositions), and select the most representative and least collinear variables. We further tested the variance inflation factor among selected variables to test the independence of the variables in the ordination space. According to *ClustOfVar*, the variables that best described environmental differences between sites were seasonality in temperature, sulphur deposition and pH (Suppl. Fig. 3). The robustness and significance of the RDAs was then tested using permutation tests.

We tested the relationships between network indices and the enviro-climatological conditions that were returned by the RDA models as most significant – temperature seasonality and sulphur deposition – using regression analyses. We grouped network indices into three categories: network composition (linkage density, species richness, partner diversity), network structure (robustness, niche specialization, modularity) and network generality (generality indices). Network indices were previously standardized to allow direct comparison among the different categories.

## Results

Mean annual temperature (MAT) was lowest in Degerö Stormyr and highest in Dosenmoor, while seasonality in temperature (TS) followed an opposite pattern (Table 1). Mean annual precipitation (MAP) varied from 710 mm in the Cena Mire to 760 mm in Dosenmoor. Seasonality in precipitation (PS) did not vary much between sites (Table 1). Total nitrogen and sulphur deposition were lowest in Degerö Stormyr. Total nitrogen deposition values in Dosenmoor were almost double as compared to those in Cena Mire. For sulphur, Cena Mire and Dosenmoor had similar deposition values (Table 1). pH in all sites was below 4.5 but differed slightly between sites (F = 4.5, *P* = 0.034) and was highest in Degerö Stormyr and lowest in the Dosenmoor (Table 1).

**Table 1.** Bioclimatic data for Degerö Stormyr, Cena Mire, and Dosenmoor. MAT = Mean annual temperature (°C), TS = Seasonality in temperature (%), MAP = Mean annual precipitation (mm), PS = Seasonality in precipitation (%); N_tot_ = Total nitrogen deposition (mg m^-2^ yr^-1^), S_tot_ = Total sulphur deposition (mg m^-2^ yr^-1^), and pH of the porewater (n=5). Different letters represent significant difference (P ≤ 0.05)

| | MAT | TS | MAP | PS | $N_{\text{tot}}$ | $S_{\text{tot}}$ | pH |
| --- | --- | --- | --- | --- | --- | --- | --- |
| Degerö Stormyr | 1.2 | 9.3 | 730 | 26 | 130 | 96 | $4.01 \pm 0.15^a$ |
| Cena Mire | 6.4 | 7.9 | 710 | 31 | 760 | 344 | $3.71 \pm 0.17^{ab}$ |
| Dosenmoor | 8.0 | 5.9 | 760 | 19 | 1260 | 6144 | $3.63 \pm 0.29^b$ |

### Plant and prokaryote community composition

Vascular plant and bryophyte species richness were highest in Cena Mire, while evenness in this site was lowest (Table 2). Detected archaeal richness did not differ among the three sites. Richness of the acrotelm archaeal communities was at least three times lower as compared to richness of the catotelm archaeal communities (Table 2). Bacterial richness, opposite to the patterns observed in the plant community, was lowest in Cena Mire, for both acrotelm and catotelm (Table 2).

**Table 2.** Plant species and prokaryote richness (OTUs) and evenness (Pielou’s J). Values are means ± standard errors. Prokaryotic richness and evenness have been described for the acrotelm and catotelm communities separately. Different letters represent significant difference (P ≤ 0.05) in richness or evenness data for the sites as tested by analysis of variance within four sub-communities (vascular plants, bryophytes, archaea and bacteria).

| Plant community |  | Richness | Evenness |  |  |
| --- | --- | --- | --- | --- | --- |
| Vascular plants | Degerö Stormyr | 6.5 ± 1.23<sup>b</sup> | 0.8 ± 0.05<sup>a</sup> |  |  |
|  | Cena Mire | 12.0 ± 1.94<sup>a</sup> | 0.2 ± 0.05<sup>b</sup> |  |  |
|  | Dosenmoor | 7.2 ± 0.40<sup>b</sup> | 0.8 ± 0.06<sup>a</sup> |  |  |
| Bryophytes | Degerö Stormyr | 1.8 ± 0.17<sup>b</sup> | 0.5 ± 0.15 |  |  |
|  | Cena Mire | 3.3 ± 0.56<sup>a</sup> | 0.5 ± 0.12 |  |  |
|  | Dosenmoor | 1.7 ± 0.21<sup>b</sup> | 0.7 ± 0.12 |  |  |
| Prokaryotic community |  | Acrotelm (oxic) |  | Catotelm (anoxic) |  |
|  |  | Richness | Evenness | Richness | Evenness |
| Archaea | Degerö Stormyr | 14 ± 2.0 | 0.8 ± 0.05 | 51 ± 4.5 | 0.4 ± 0.06 |
|  | Cena Mire | 30 ± 13.3 | 0.6 ± 0.08 | 74 ± 7.0 | 0.4 ± 0.05 |
|  | Dosenmoor | 38 ± 14.2 | 0.7 ± 0.05 | 84 ± 11.9 | 0.5 ± 0.06 |
| Bacteria | Degerö Stormyr | 1367 ± 95.0<sup>a</sup> | 0.8 ± 0.00 | 1511 ± 99.4<sup>a</sup> | 0.8 ± 0.01<sup>a</sup> |
|  | Cena Mire | 1231 ± 109.0<sup>b</sup> | 0.8 ± 0.02 | 838 ± 105.9<sup>b</sup> | 0.7 ± 0.04<sup>b</sup> |
|  | Dosenmoor | 1615 ± 88.3<sup>a</sup> | 0.8 ± 0.01 | 1246 ± 139.2<sup>a</sup> | 0.7 ± 0.01<sup>a</sup> |

Redundancy analysis (RDA) showed that all communities (plants, acrotelm prokaryotes, and catotelm prokaryotes) well separated in the ordination space (Suppl. Fig. 4). RDA analysis explained 28% (vascular plants), 7.2% (prokaryote, acrotelm) and 19% (prokaryote, catotelm) of the variance (adjusted *R*^2^) of each community. Sulphur deposition and temperature seasonality both explained much of the differences in community structure for all three communities (*P* ≤ 0.05), while pH was not significant. Notably, the three communities showed similar patterns in their species-environment relationships. Communities in Degerö Stormyr were related to high seasonality in temperature, and communities in Dosenmoor were related to high sulphur deposition (Suppl. Fig. 4).

Across sites, and pooled for the acro- and catotelm communities, bacterial to archaeal ratios exceeded 3:1 (Fig. 2 a). Acidobacteria (35%), Proteobacteria (29%), Verrucomicrobia (13%), and Actinobacteria (9%) were the most abundant bacterial phyla across sites (Fig. 2 b). Relative abundances in bacterial phyla were similar across sites. Composition of the archaeal community, on the other hand, seemed site-specific (Fig. 2 c). In Degerö Stormyr, methanogens of the order Methanomicrobiales (50%) and Methanobacteriales (37%) were the dominant archaeal orders.

**Figure 2.**
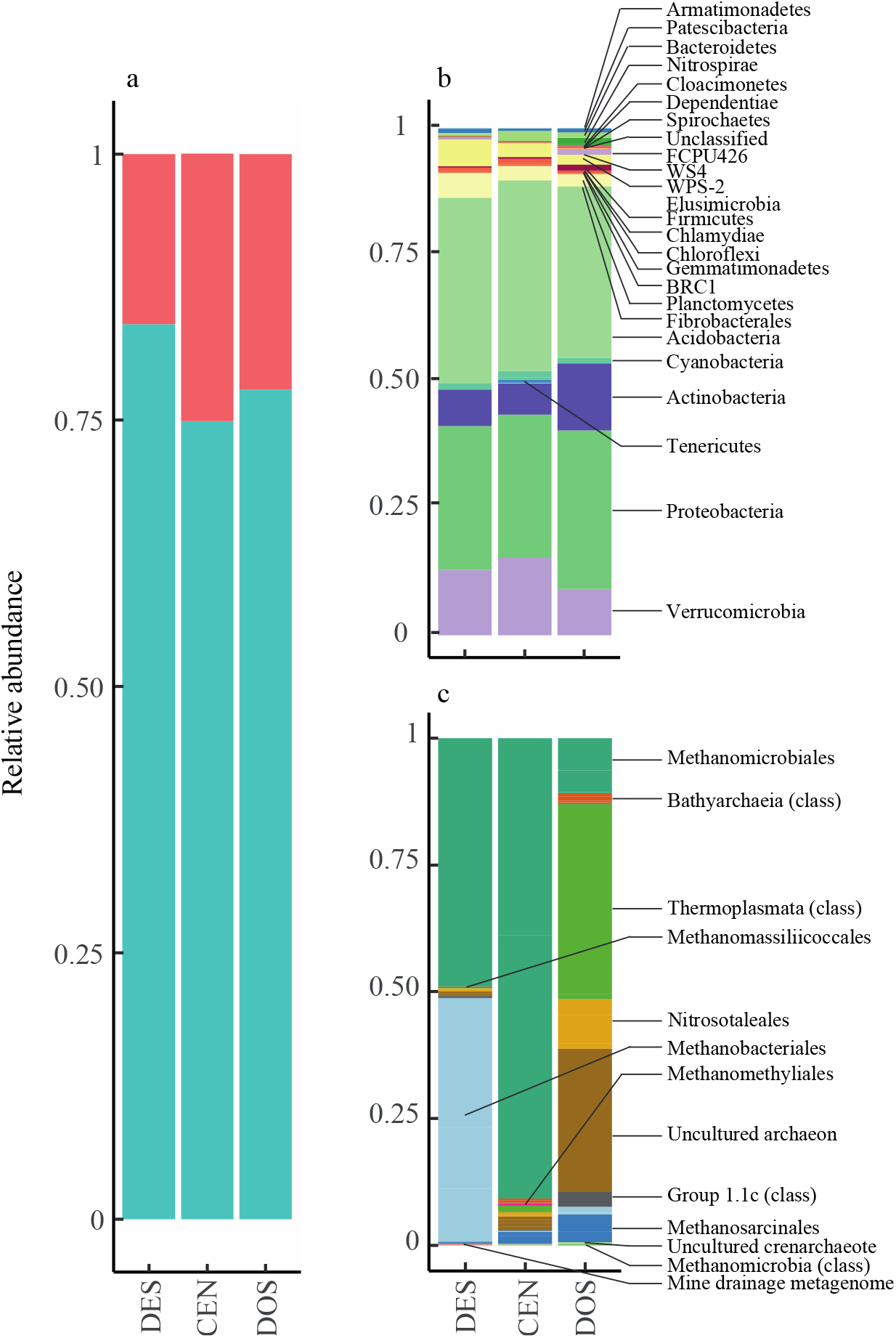
Comprehensive view of the sequence content of peat soil prokaryotic libraries per site: DES = Degerö Stormyr, CEN = Cena Mire, DOS = Dosenmoor. Segments that compose each bar represent the total number of sequences normalised to the total number of sequences in the libraries per site. a) View of the relative abundance of bacteria (blue) and archaea (red) in the site-specific communities. b) View of the relative abundance of bacterial ‘species’ in the respective site-specific communities. Each segment represents identified bacterial phyla. c) View of the relative abundance of archaeal ‘species’ in the respective site-specific communities. Each bar segment represents identified archaeal orders.

Archaeal communities in Cena Mire were dominated by one single order, Methanomicrobiales (83%). In Dosenmoor, Thermoplasmata (39%) and an uncultured archaeon (28%) likely associated to the phyla Thaumarchaeota and Crenarchaeota were most dominant. Of the classified orders in this site, Nitrosotaleales (30%), Methanomicrobiales (22%) and Methanosarcinales (19%) had highest prevalence.

### Network structure

The number of plant-prokaryote pairs in bipartite networks differed between the three sites (site effect, *F_1,2_* = 65, *P* = 0.01; site x depth, *F_1,2_* = 2.2, *P* = 0.27), with > 70% fewer interactions in Cena Mire and Dosenmoor as compared to Degerö Stormyr (Fig. 3). A lower number of bipartite interactions results from lower numbers of both positive and negative associations (Fig. 3).

**Figure 3.**
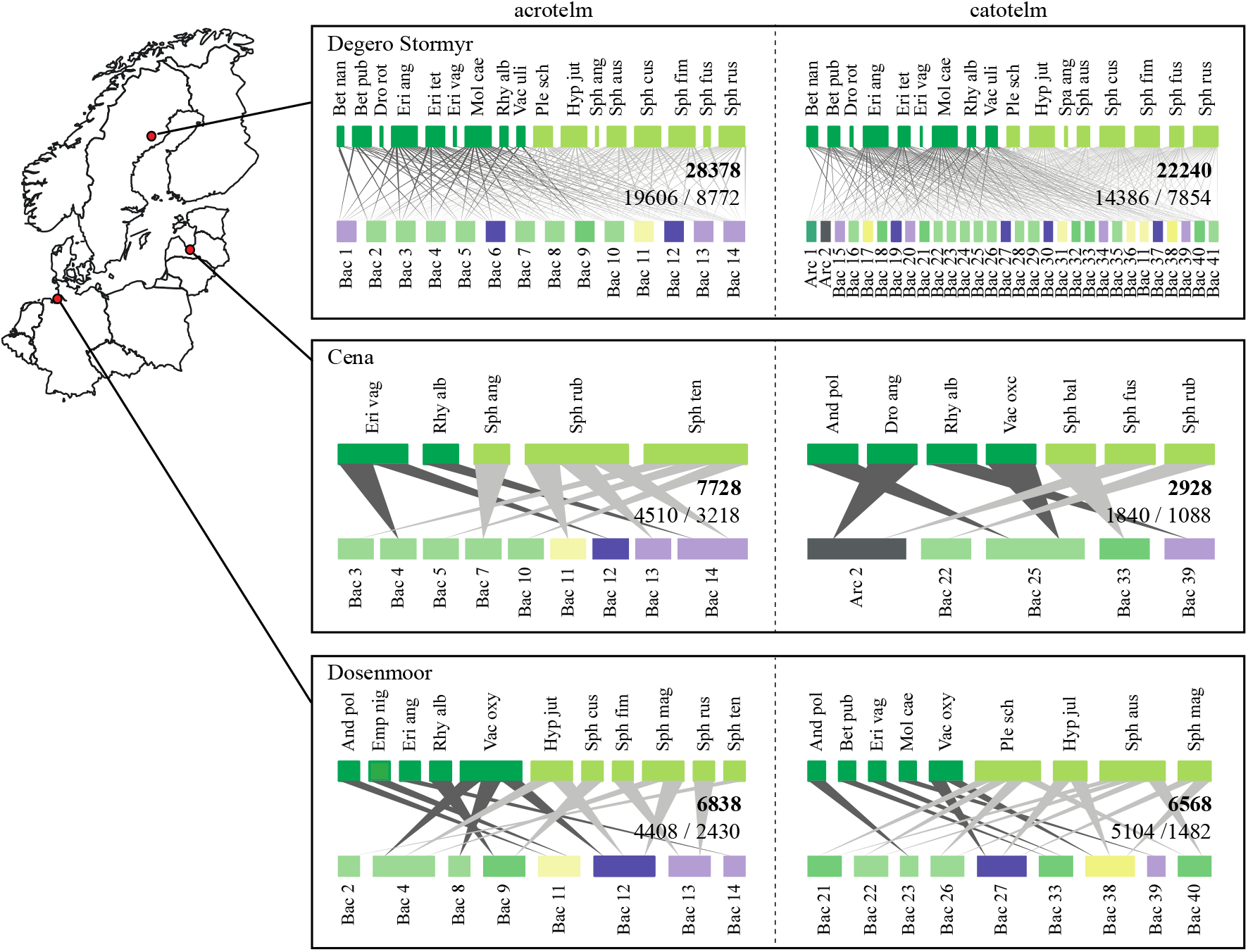
Bipartite network visualisation for the three European peatlands. Indicated are interactions between the plant community and acrotelm prokaryotic community (left) and between the plant community and catotelm prokaryotic community below (right). Plants are visualised at the top of the networks and coloured dark and light green for vascular plants and bryophytes, respectively. Prokaryotes are illustrated on the bottom of the networks and follow the colour scheme as in Fig.1. A complementary list of prokaryotes and their classification can be found in Suppl. Table 1. Size of the bars scales with relative abundance in the respective communities. For all sites, networks are filtered to only visualise interactions with ‘keystone’ prokaryotes, defined as those that had > 5 interactions in Degerö Stormyr. Interaction lines represent Spearman’s rank correlations > 0.6. Dark grey interactions indicate vascular plant-prokaryote interactions, light grey indicates bryophyte-prokaryote interactions. Inset numbers indicate the full number of bi-partite (bold) interactions, further broken down in the number of positive / negative interactions.

In the vascular plant community, only *Rhynchospora alba* could be identified to play a major role in all plant-prokaryote networks, except in the Dosenmoor catotelm network (Fig. 3). In the moss community, no such key species seems present (Fig. 3). OTUs classified as belonging to the families Acidobacteriaceae (Bac 4), WD2101 soil group (Bac 11), Acidothermaceae (Bac 12), Pedosphaeraceae (Bac 13), and Methylacidiphilaceae (Bac 14) were present in all three networks of the acrotelm prokaryotic communities (Fig. 3, Suppl. Table 1). In the catotelm prokaryotic communities, bacteria in the families Koribacteraceae (Bac 22), Acetobacteraceae (Bac 33), and Pedosphaeraceae (Bac 34) were important network species (Fig. 3, Suppl. Table 1).

For all sites, β-diversity for bipartite (plant-prokaryote) networks (BDN) exceeded β-diversity for community composition (BD), indicating that network turnover is higher than species turnover between sites (*P* ≤ 0.001; Fig. 4). Indeed, network compositions (i.e. pairwise comparison between networks) between sites were more dissimilar (acrotelm, BDN = 0.91; catotelm, BDN = 0.89) than species compositions (i.e. pairwise comparison between communities) between sites (acrotelm, BD = 0.32; catotelm, BD= 0.31). Turnover in species composition between sites, β-diversity (BD), was equally driven by species and local contributions (BD_S_ and BD_L_). Network β-diversity (BDN) was mostly driven by turnover in interactions (both direction and number) between species that are common in the three sites (BDN_CS_), rather than by novel interactions driven by changes in the community composition (BDN_NS_; Fig. 4).

**Figure 4.**
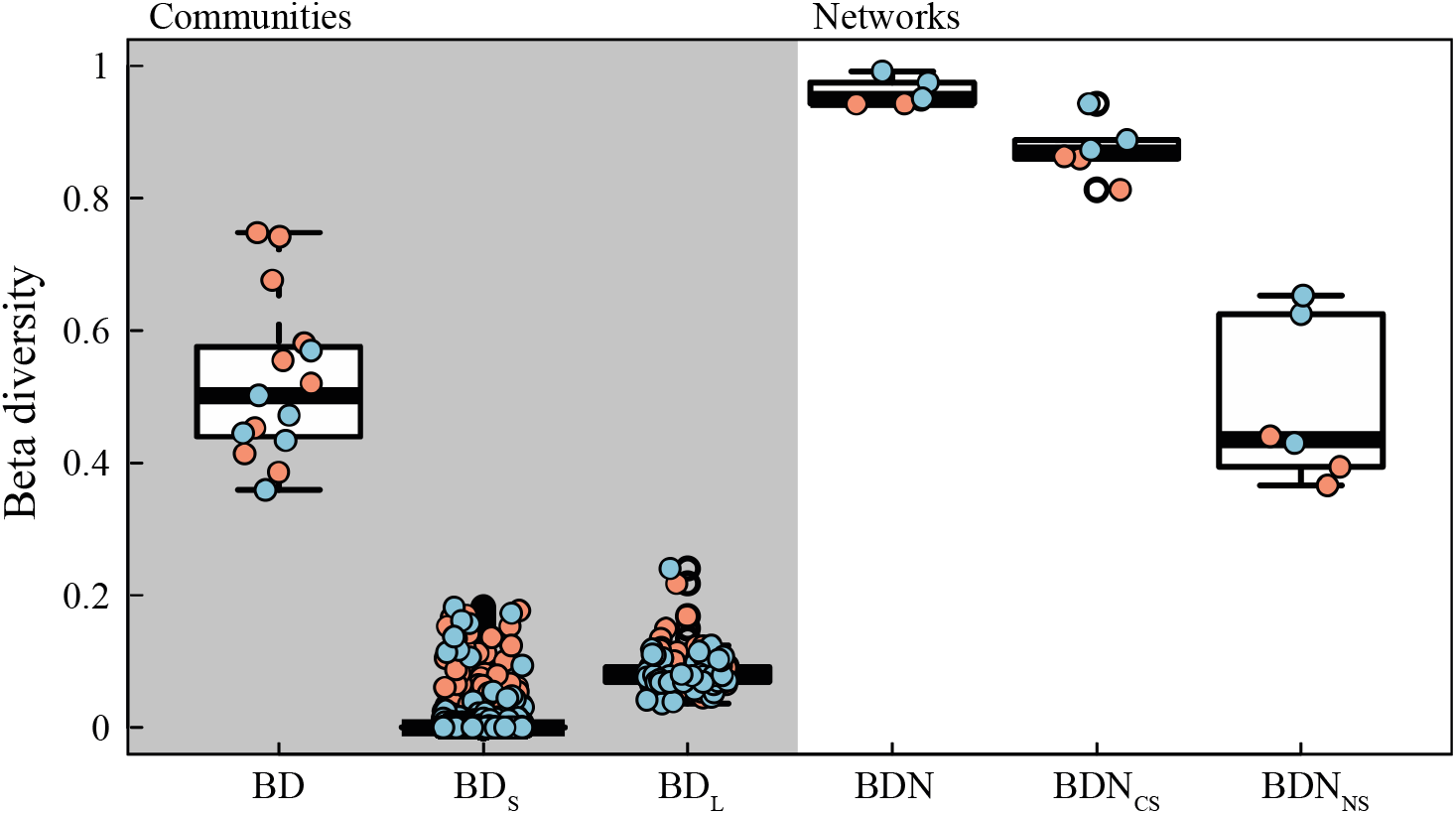
Turnover in species composition (species β-diversity) and association (network β-diversity) and their respective components across three peatlands (see methods). Red circles indicate plant and microbial communities in the acrotelm. Blue circles indicate plant and microbial communities in the catotelm.

Differences in the number of bi-partite interactions between sites seem to relate to differences in the numbers of effective partners between the respective communities, which were always highest in Degerö Stormyr (acrotelm: *P* < 0.001; catotelm: *P* < 0.001; Fig. 5 a, b). This is further supported by differences in diversity in plant-prokaryotic partners, which were higher in Degerö Stormyr as compared to Cena Mire and Dosenmoor (acrotelm: *P* < 0.001; catotelm: *P* < 0.001, Fig 4 c, d). Moreover, the number, but also diversity, of prokaryotic OTUs that are associated (i.e. effective partners) to specific plant species (top-down links) were higher than the number of plant species that associate with a specific OTU in the prokaryote community (bottom up links). Essentially this means that plant species link to a higher number of prokaryotic species, than prokaryotes to plants. Despite the relative low number of links of prokaryotes with plants, we found a larger specificity of these links, as compared to the specificity of plants in their association with prokaryotes (acrotelm: *P* < 0.001; catotelm: *P* < 0.001). Partner specificity was lowest in Degerö Stormyr for both levels of organisation in the bi-partite networks (acrotelm: *P* < 0.001; catotelm: *P* < 0.05, Fig. 5 e, f).

**Figure 5.**
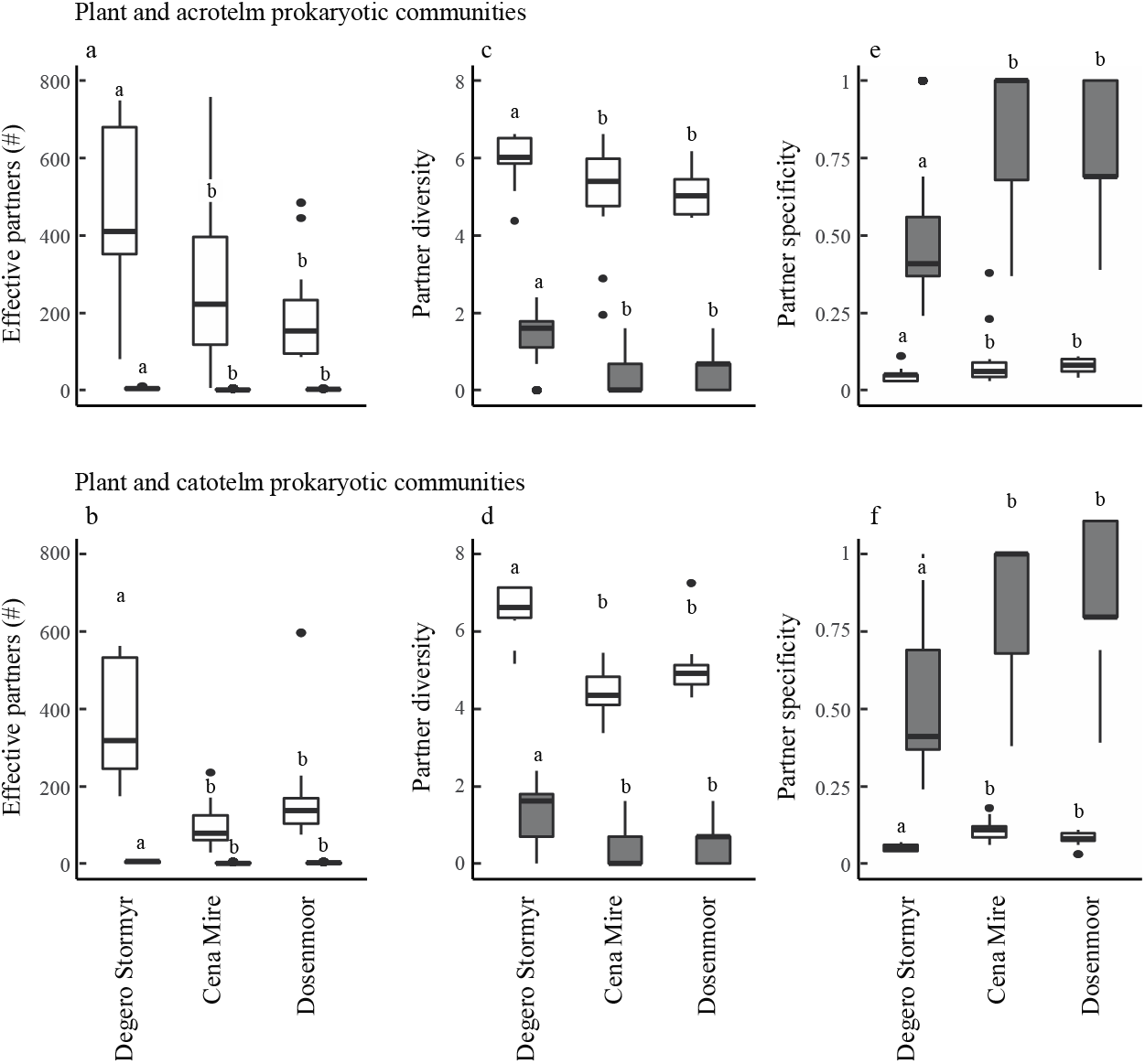
Boxplots of the species associations and network properties for the acrotelm (a, c, e) and catotelm (b, d, f) plant-prokaryote interactions. The number of specific partners (a, b), partner diversity (c, d) and partner specificity (e, f) have been calculated for plants (white boxplots: top down links; Suppl. Fig. 2) and prokaryotes (grey boxplots: bottom up links; Suppl. Fig. 2). Horizontal solid lines indicate median values. Different letters indicate differenced at the P ≤ 0.05 level, based on analysis of variance using mean values.

At the network level, the overall linkage density (LD), generality (G), partner diversity (PD) and species richness (SR) in the bipartite networks was higher in Degerö Stormyr as compared to Cena Mire and Dosenmoor; a pattern found in both acrotelm and catotelm bipartite networks (Suppl. Fig. 5). The bipartite networks, acro- and catotelm, in Degerö Stormyr were strongly sensitive to the loss of prokaryotes (low robustness) but not to the loss of plants. Oppositely, bipartite networks in Cena Mire and Dosenmoor were more sensitive to the loss of plants as compared to prokaryote loss (Suppl. Fig. 5). As compared to Degerö Stormyr, networks in Cena Mire and Dosenmoor show a high degree in niche specialization, especially in the plant communities (Suppl. Fig. 5). These differences in network indices between the sites were driven mostly by temperature seasonality and sulphur deposition in a lesser extent (Fig. 6). While compositional network indices (linkage density, partner diversity, number of effective partners) and generality indices increased with increasing temperature seasonality, structural network indices (robustness, niche extinctions, modularity) decreased. Opposite patterns were found with increasing sulphur deposition (Fig. 6). These results indicate that plant-microbial interactions change in complexity, specificity and evenness in the distribution of species interactions with changes in environmental conditions, and that these patterns are likely driven by the turnover of species between peatland sites.

**Figure 6.**
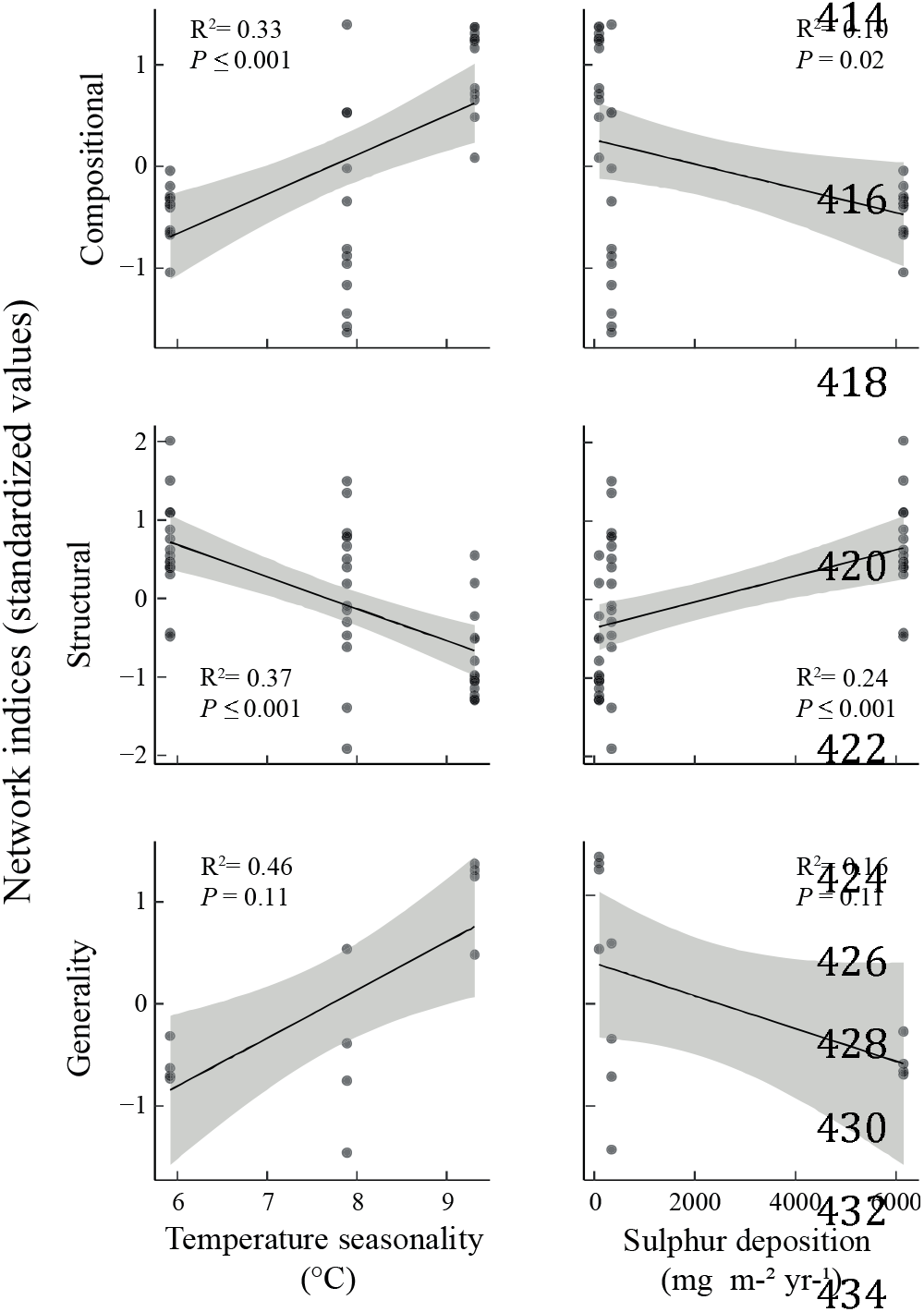
Linear regressions between the key enviro-climatological variables–total nitrogen deposition and temperature seasonality– and network indices. Network indices were grouped into three categories, network composition (linkage density, species richness, partner diversity), network structure (robustness, niche specialization, modularity) and network generality (generality indices). The grey area represents the 95% confidence interval of the mean. Data on specific network indices can be found in Suppl. Fig. 5.

## Discussion

Using data on plant and prokaryotic community composition in three *Sphagnum-dominated* peatlands, we evidence that plant-prokaryotic networks are site-specific. As these peatlands are situated in three distinct environmental zones (Metzger et al. 2005) our data suggest that these networks are vulnerable to the effects of enviro-climatological change. Notably, our work indicates that turnover in plant-microbe links outpaces turnover in biotic composition of the plant and microbial communities that comprise the plant-microbe networks. This suggests a disproportionate rewiring of plant-microbe networks along an enviro-climatological gradient, as hypothesized in scenario iv (Fig. 1). Hence, a turnover in the plant community as shown in earlier research (Gunnarsson et al. 2002, Robroek et al. 2017, Pinceloup et al. 2020) will lead to complex changes in plant-microbe interactions that could affect ecosystem processes in the longer term (Morriën et al. 2017).

### Turnover in community composition

In line with earlier work where it was suggested that changes in enviro-climatological conditions in peatland ecosystems leads to a significant turnover in the composition of the plant community (Robroek et al. 2017), plant community composition was highly distinctive between sites. Moreover, our study confirms previous suggestions that enviro-climatological conditions are important drivers that underlie the composition of the vegetation, and that changes in these drivers result in alterations of the plant community (Gunnarsson et al. 2002, Pinceloup et al. 2020) and associated biotic interactions (Wiedermann et al. 2007). Indeed, we note that site-specificity of plant communities extends to the prokaryotic community, a pattern that is likely explained by the strong links between plant and microbial communities (Chronakova et al. 2019, Ivanova et al. 2020). Despite the variation in prokaryotic community composition between sites, the archaea to bacteria ratio remains strikingly stable, suggesting that both microbial groups are equally affected by alterations in enviro-climatological conditions.

Bacterial community composition was remarkably similar between sites. Archaeal communities, on the other hand, were highly dissimilar between the three peatland sites. Bacteria in the genera *Occallatibacter, Acidothermus, Verrucomicrobium*, bacteria in the Phycisphaera-related group WD2101 (yet uncultured Planctomycetes of the order Tedisphaerales) and bacteria in the family Methylacidiphilaceae (phylum Verrucomicrobia) occurred in all three acrotelm networks. These genera have been previously found in soils, and have been characterised as strictly aerobic, moderately acidophilic, and chemoorganotrophic mesophiles. Methylacidophilaceae include methane-oxidizing genera, but there is no evidence that the members of this family present in peatlands are involved in methane cycling (Ivanova et al. 2020). This underlies the fact that it can be difficult to confidently predict the function of uncultivated bacterial species. The bacterium “*Candidatus* Koribacter”, a bacterium in the family Acidobacteriaceae, and a bacterium in the family Pedosphaeraceae were important network species in catotelm prokaryotic communities. These results mirror earlier findings, where bacterial community composition was reported to be rather stable under experimental warming (Weedon et al. 2017). The archaeal communities in Degerö Stormyr and Cena mire were dominated by methanogens. In these sites, biogeochemical drivers likely promoted a classical anaerobic degradation of organic matter to methane in the absence of other terminal electron acceptors. The dominating methanogens, Methanobacteriales and Methanomicrobiales, characteristic for acidic peatlands, are hydrogenotrophic and form methane from hydrogen/CO_2_ and/or formate (Kotsyurbenko et al. 2007, Martí et al. 2015). Interestingly, the Methanomicrobiales family *Methanoregulaceae* identified in the present study dominated the methanogen population in a comparable set of peatlands along a nitrogen deposition gradient (Martí et al. 2015). Reductions in the relative abundance of methanogens in Dosenmoor may be the result of higher deposition levels of nitrogen and sulphur. The presence of oxidized forms of these elements, may favour nitrogen oxide and sulphate reduction, resulting in increased competition for methanogens. This is supported by the higher relative abundance of the phyla Nitrosotaleales and Thaumarchaeota (Suppl. Table 1), which are known ammonia-oxidizing archaea in acidic environments (Lehtovirta-Morley et al. 2011, Lin et al. 2015), in the Dosenmoor. Furthermore, Thermoplasmata – highly abundant in the Dosenmoor – are suggested to be involved in the transformation of sulphite and possibly organosulphonates (Lin et al. 2015). Given the high sulphur deposition in the Dosenmoor, these compounds prospectively are abundant. Here, it should be stressed that relative abundance does not necessarily correspond to the actual abundance and/or the physiological activity by the microorganisms. To exemplify, increased methanogen activity has been shown in peatlands with high nitrogen deposition and low relative abundance in methanogens (Martí et al. 2019a). Hence, the main part of the organic matter degradation likely follows the methanogenic route, also at the Dosenmoor. In this context it should be noted that the relative abundance might discriminate against prokaryotes at very low abundance, which may have very high activity and thus process biogeochemical reactions at high speed as exemplified for sulphur in peat soil (cf. Pester et al. 2010, 2012)

### Turnover in plant-microbe network composition

Beyond differences in community composition, our work shows that plant-microbe network structure differs between sites. Notably, and in line with scenario iv in our hypotheses, turnover of plant-microbe interactions is disproportionate when compared to the turnover in plant and prokaryote communities. These results would suggest that changes in enviro-climatological conditions are key in driving the observed rewiring of plant-prokaryote interactions. Interestingly, the highest numbers of effective partners and partner diversity were found in the plant community. While prokaryotes have significantly lower numbers of plant partners, their associations with plant species tend to be more specific. The observation that diversity and numbers in top-down interactions (i.e. number of interactions plant species have with prokaryotes) exceeds diversity and numbers in bottom-up interactions (interactions prokaryotes have with species in the plant community) can simply be attributed to the fact that species richness in the prokaryotic community exceeds plant species richness by a couple of orders of magnitude, which by definition leads to higher diversity of plant-associated prokaryotes. Our analysis, however also suggests that species in the soil prokaryotic community in peatlands form specific links to species in the plant community (cf. Bragina et al. 2012, Hamard et al. 2019). Three non-exclusive scenarios can explain the patterns we observe. First, a particular plant species provides resources or niche space for a wide array of microbes. Second, a particular plant species under particular environmental conditions produce and release specific metabolites that can attract, deter or kill targeted microbes. Third, these microbes are highly specific to a particular plant (Suppl. Fig. 2). Taken together, our findings indicate that plants are important drivers for plant-prokaryote interactions and is in line with the general idea that the plant community has a direct effect on the prokaryotic species occurrences and community composition and activity (Fisk et al. 2003, Wardle et al. 2004, Robroek et al. 2015, Yavitt & Williams 2015), but the opposite is not necessarily true.

Network size and complexity differ between sites and decrease from the most northern (Degerö Stormyr) to the most southern (Dosenmoor) site. These findings are in line with earlier findings where, as a response to warming, networks were observed to become less complex (Galiana et al. 2014). Wei et al. (2013) have connected nutrient enrichment to a decoupling of grassland plant-soil-microbe interactions and attribute this to differences in responses in the composition of plant and microbial communities to changes in nutrient load. Hence, the observed decrease in network complexity along the enviro-climatologic gradients in our study may indicate that plant-microbe associations, and perhaps even feedbacks, go down with increased nutrient deposition and decreased temperature seasonality.

A faster loss of network interactions as compared to species loss has earlier been suggested to have major implications on ecosystem processes and services (Valiente-Banuet et al. 2014). Indeed, increased network complexity has been connected to enhanced efficiency in soil carbon uptake, that is increased C uptake by the soil microbial communities without substantial changes in microbial biomass, even without major changes in the composition of plant communities (Morriën et al. 2017). Our data suggest that seasonality in temperature and nutrient deposition modulate network indices that are related to composition, structure and generality of plant-microbe networks. In other words, our findings indicate that decreased seasonality and increased nutrient deposition, which reflect a north-south gradient in our study, leads to decreases in network indices related to composition and generality, but simultaneously increases structure related indices. A number of plant-prokaryotic associations that were identified in bi-partite networks in Degerö Stormyr were persistent in the other two sites despite differences in enviro-climatological different. These results would suggest a certain resistance and robustness of plant-prokaryotic associations when facing radical environmental change (Mandakovic et al 2018). Furthermore, while networks become less diverse and complex while moving from the northern to the southern end of our enviro-climatologic gradient, network structure becomes less prone to species loss. Essentially this means that we observe an increased level of specialization in ecological networks, potentially caused by local adaptations that favour network specialization (Dalsgaard et al. 2011, 2013) under harsh conditions (e.g.. increased nutrient load) for peatland ecosystems. Such increase in specialization indicates an increased level of nestedness and would in the long term weaken network stability to changing enviro-climatological conditions (de Vries et al. 2018).

### Network rewiring – new connections between common species

As correlation networks along environmental gradients between vegetation and microbial communities may not necessarily be causal, care should be taken in drawing firm conclusions from our data. Additionally, space-for-time substitutions, as used in this study, have been reported to show a greater magnitude of response in community turnover than climate manipulations or long-term monitoring studies (Elmendorf et al. 2015). Space-for-time approaches nevertheless are extremely useful for gauging the direction of change of communities to decadal scales in enviro-climatological change (Elmendorf et al. 2015). Despite these caveats, our findings show that while differences in environmental properties, such as temperature and nutrient deposition, between sites markedly relate to differences in the composition of plant and prokaryote, acro- and catotelm, communities, turnover in plant-prokaryote network composition exceeds species turnover in either one of both communities. Specifically, we show that a reshuffling of plant and microbial communities, most likely driven by enviro-climatological changes, has resulted in a significant rewiring of plant-microbial interactions (scenario 4; Fig 1). This rewiring was not just the result of changes in spatial patterns of diversity across plant and microbe trophic levels (cf. Galiana et al. 2019). The rewiring between the peatland plant-microbe networks that we have assessed was to a large degree caused by the creation of novel associations between species that are common and shared across the sites. Hence, turnover in network composition is largely driven by novel interactions between plants and microbes that are common across networks and to a lower extent by species new to a metacommunity due to species turnover. These results would suggest that changes in environmental conditions can lead to network rewiring, without clear and strong effects in plant or microbial community per se. We propose that the next step is to link changes in plant-microbe network structure to peatland carbon cycling and to identify key plant and microbial groups that underpin the carbon uptake function of northern peatlands.

## Data and materials availability

All data that support this manuscript are available through DANS, the Netherlands institute for permanent access to digital research resources (https://dans.knaw.nl/en). Nucleotide sequence accession numbers have been deposited in the European Nucleotide Archive (ENA) at EMBL-EBI under accession number PRJEB36623.

## Declarations

### Acknowledgments –

We thank the Danish National High-Throughput DNA-Sequencing Centre (Copenhagen, Denmark) for using their sequencing facilities. We also thank the members of the PEATBOG project for valuable input in the planning phase of this project, and the landowners for field access. This paper is dedicated to late Dr Richard Payne, who tragically lost his life while attempting to climb Peak 6477 in the Indian Himalayas.

### Author contributions –

BJMR and BHS initiated the data collection. BJMR and MM led the field sampling. VEJ, MM and BJMR analysed all data, with help from MGD, AJV and BHS, and led the writing of the manuscript to which all co-authors contributed. The first and second author contributed equally to this paper.

## Supplementary Tables and Figures

**Supplementary Table 1.**
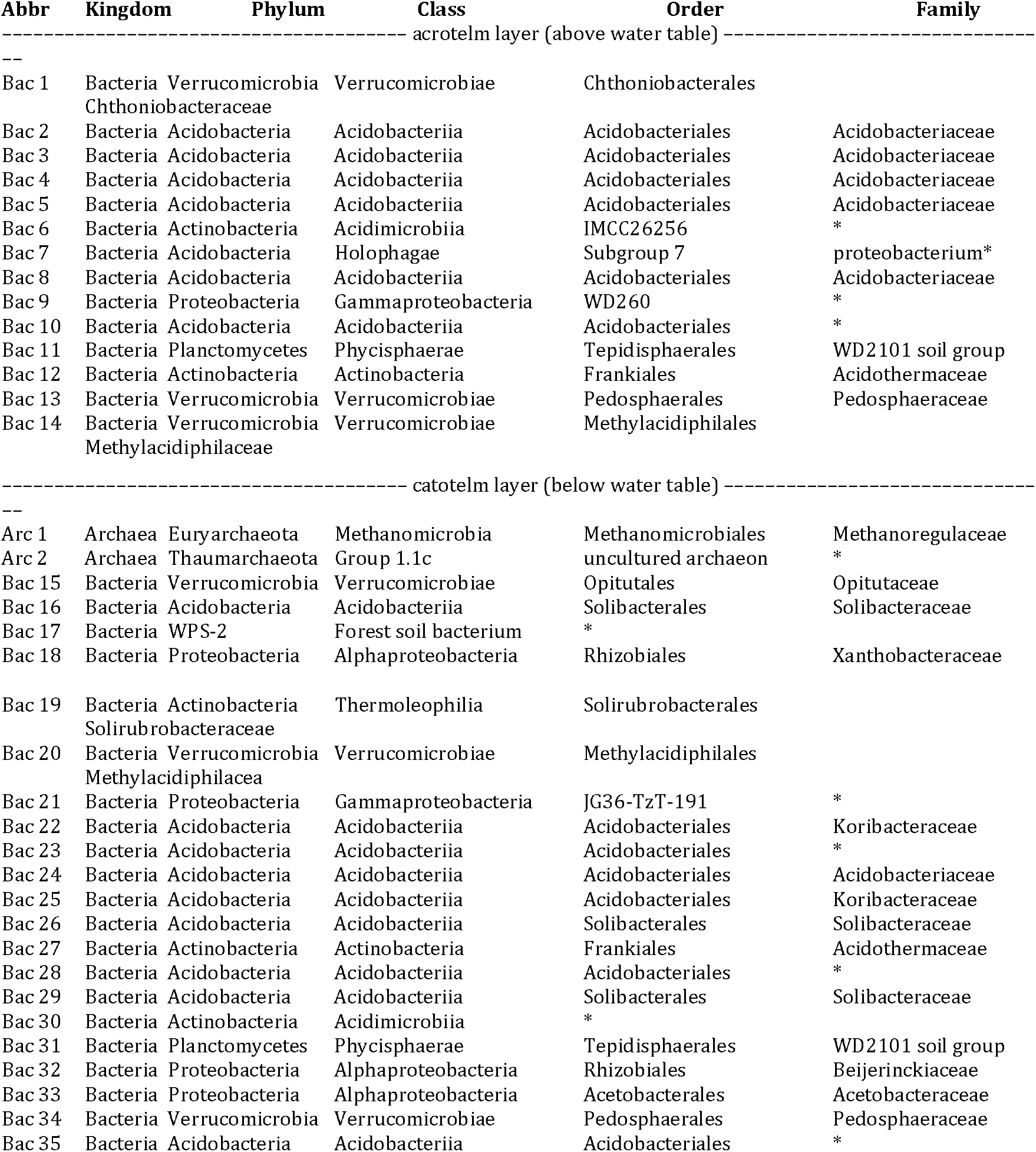
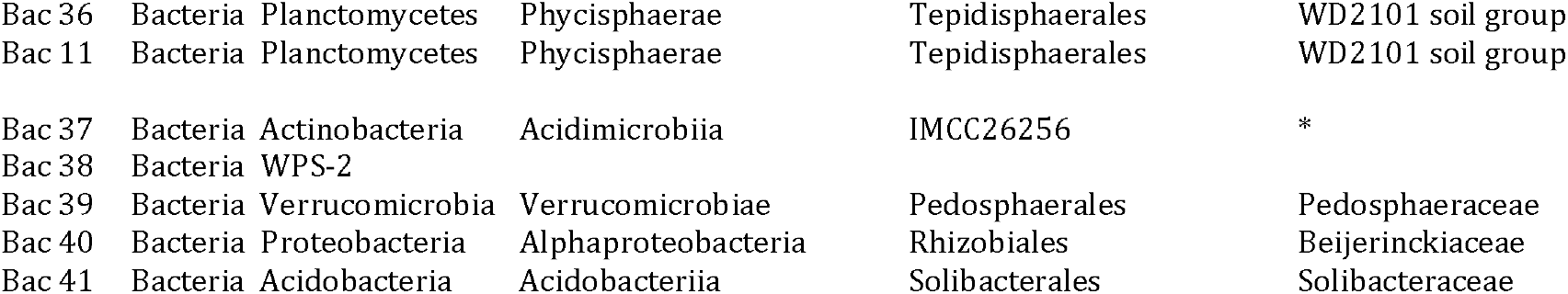
Key prokaiyotes in the networks (OTUs with > 8 interactions], * indicate uncultured prokaryotes.

**Supplementary Figure 1.**
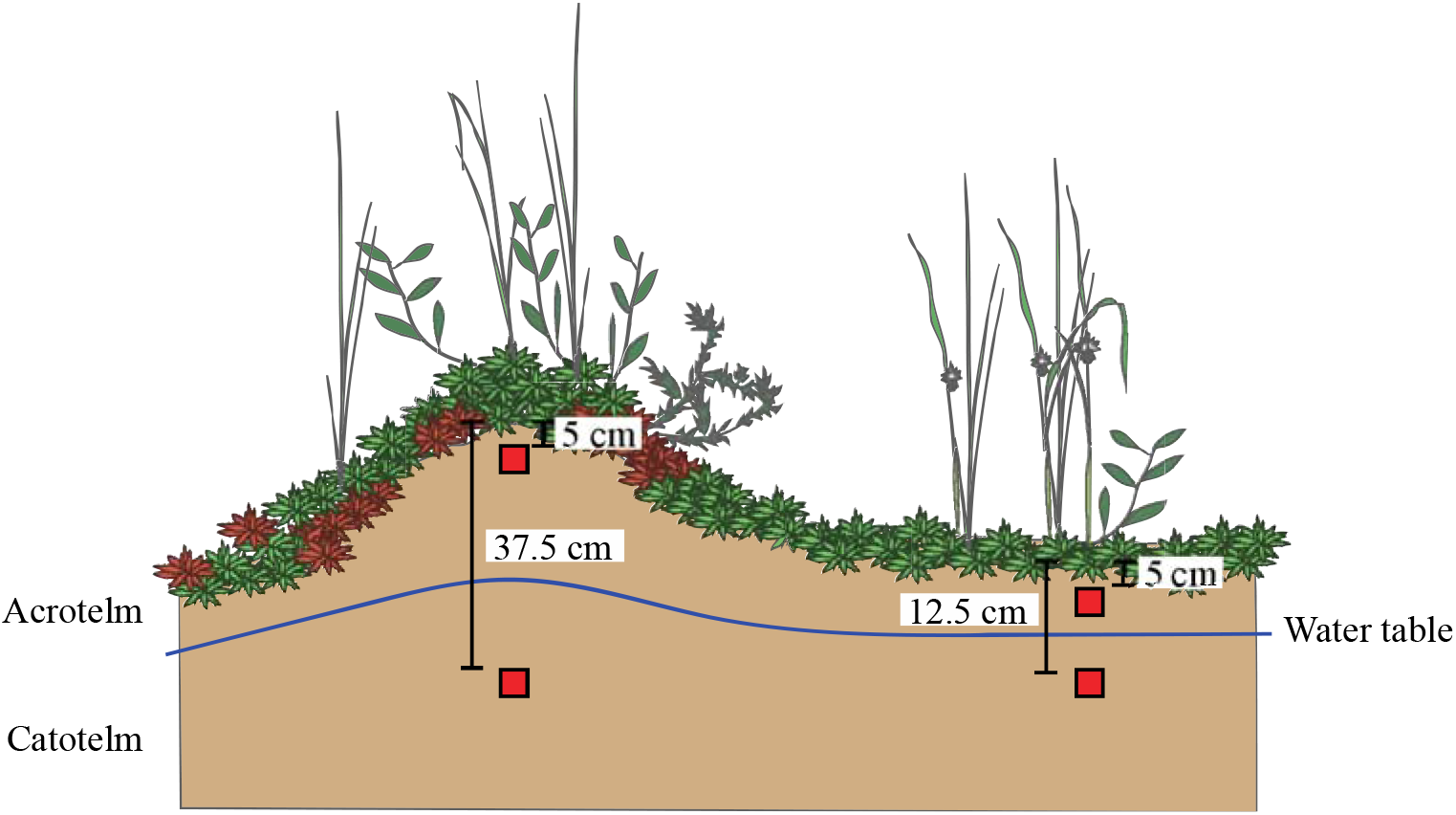
Schematic cross-cut of a typical *Sphagnum*-dominated peatland with a distinct lawn and hummock (i.e. peat mound) microhabitat pattern. In the both microhabitats, samples were extracted from the acrotelm and the catotelm peat. Acrotelm peat samples were takes 5 cm below the peat surface in both microhabitats. Catotelm peat samples were taken 12.5 cm and 37.5 cm below the peatland surface in the lawn and hummock microhabitats, respectively.

**Supplementary Figure 2.**
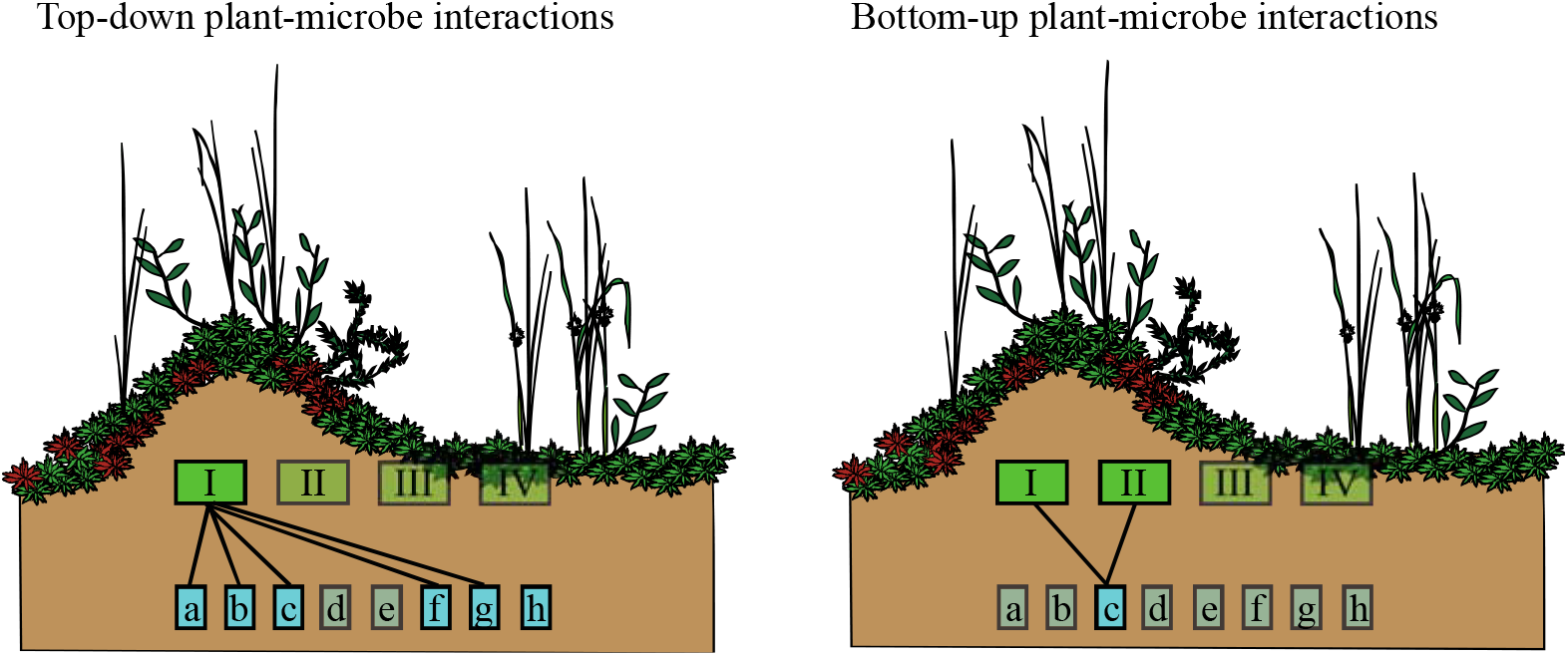
Plant-microbe bipartite network interactions. Each plant (I) in the plant community (I-IV) interacts with a number of prokaryotic ‘species’ (a,b,c,f,g & h). Hence, top-down interactions (left) can be quantified for all plant-microbe interactions in terms of network attributes (see main text). In the same network (right), a prokaryotic ‘species’ (c) can interact with species in the plant community (I & II). Again, these bottom-up interactions can be quantified for all microbe-plant interactions in terms of network attributes. It is important to note that top-down and bottom-up network attributes can differ significantly (see Fig. 2, and main text) as the number interactions of plant species with microbial ‘species’ does not necessarily parallel the number of interactions of microbial species with plant species.

**Supplementary Figure 3.**
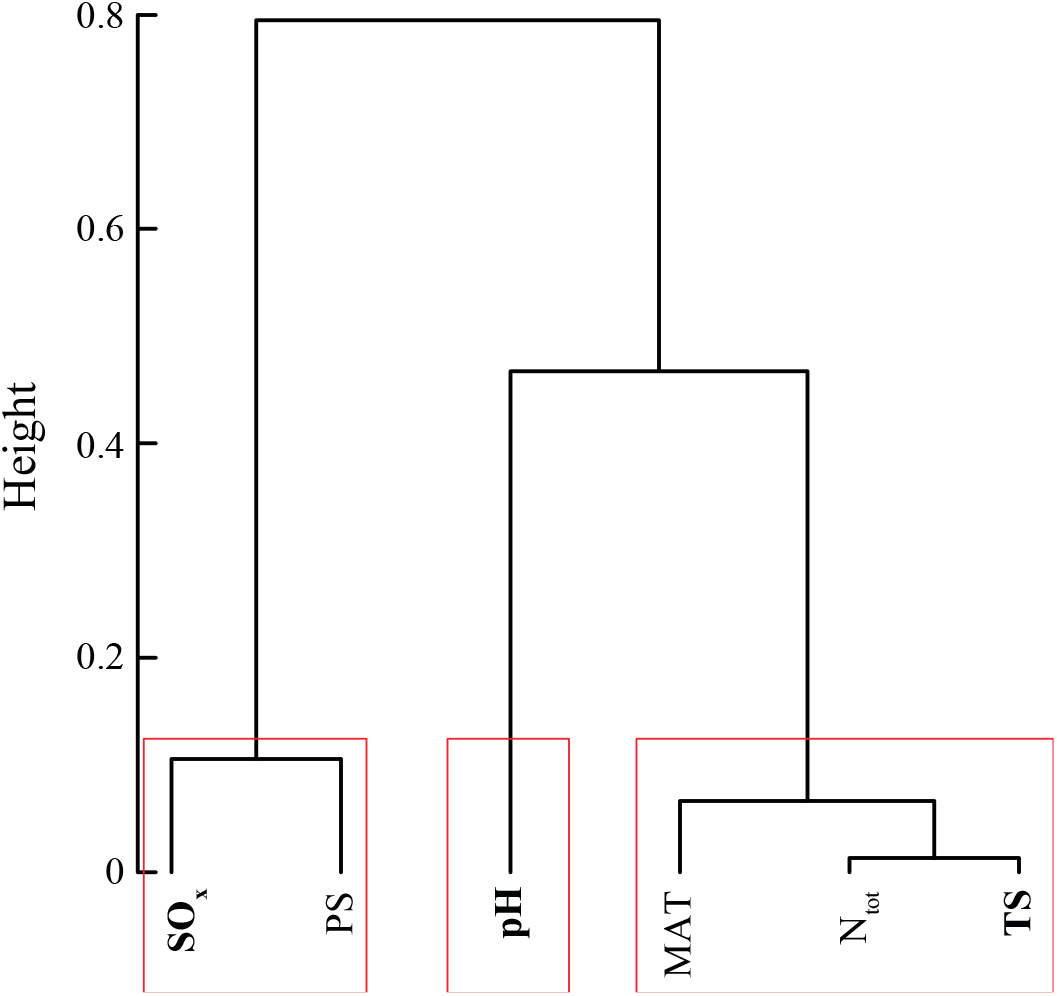
Enviro-climatological variable reduction using the *ClustOfVar* function (Chavent et al. 2012). Following *ClustOfVar*, three variables were selected (bold) from the different clusters (in red): SO_x_, pH and TS.

**Supplementary Figure 4.**
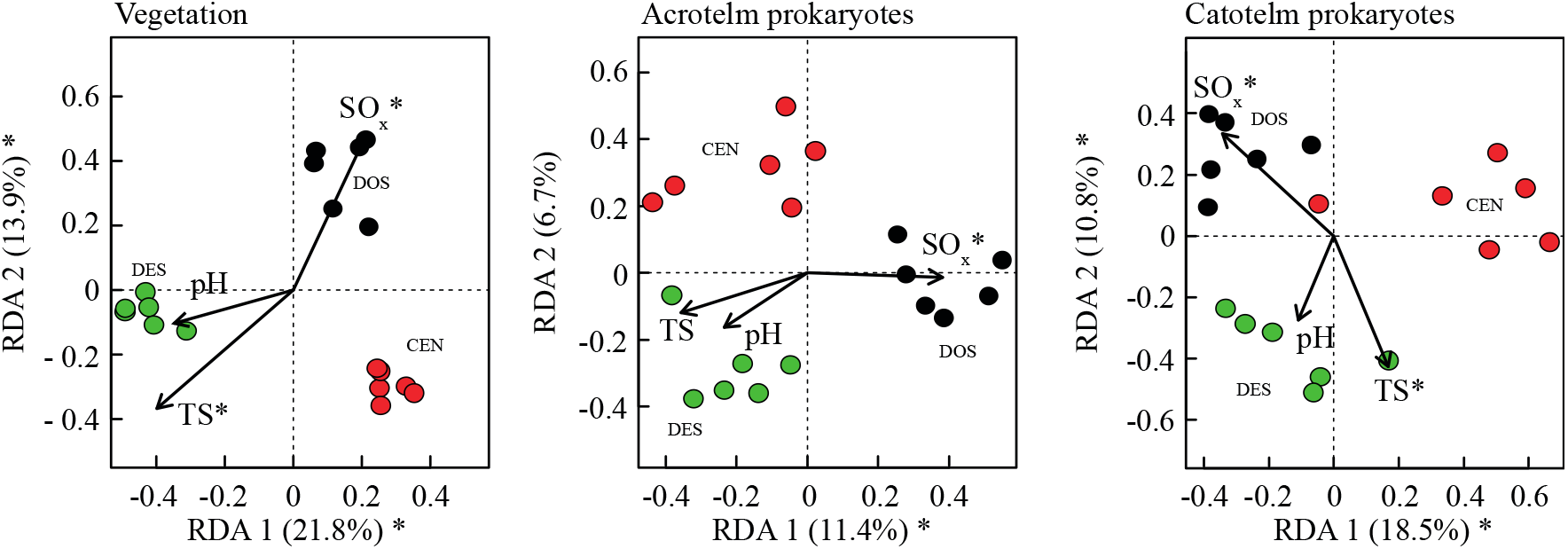
Redundancy analyses (RDA; axes 1 and 2) biplot of site scores for the plant species (left), prokaryotic OTUs sampled from the acrotelm peat (centre), and prokaryotic OTUs sampled from the catotelm peat (right). RDA scores are constrained by data on pH, seasonality of temperature (TS) and sulphur deposition (N). * indicates enviro-climatological variable that were returned as significant (*P* ≤ 0.05) by the RDA models.

**Supplementary Figure 5.**
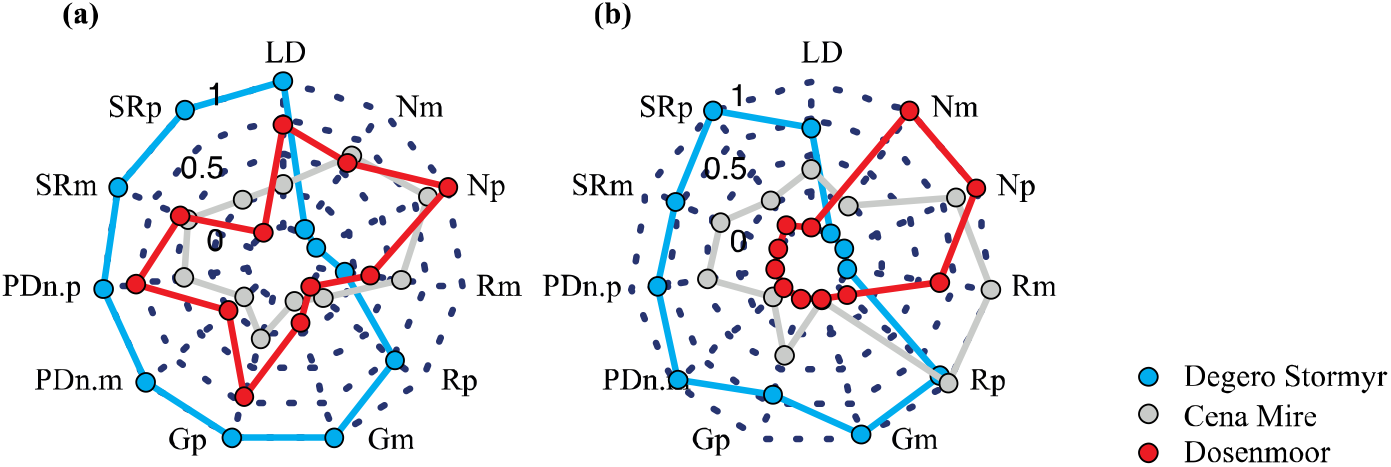
Network topology for the (a) plant–acrotelm prokaryote and (b) plant–catotelm prokaryote bi-partite networks, calculated at network level. LD = Linkage density (LD), Gp = Generality of plant links with prokaryotes. Gm = Generality of prokaryote links with plants, R = Robustness, PDn = Partner diversity in the networks, SR = Species richness (SR), N= Niche specialization (N). The indices G, R, PDn SR and N were calculated separately for the two levels –plants (p) and prokaryotes (m)–in the bipartite networks.

**Rpackages**

